# 3D Genome Analysis Identifies Enhancer Hijacking Mechanism for High-Risk Factors in Human T-Lineage Acute Lymphoblastic Leukemia

**DOI:** 10.1101/2020.03.11.988279

**Authors:** Lu Yang, Fengling Chen, Haichuan Zhu, Yang Chen, Bingjie Dong, Minglei Shi, Weitao Wang, Qian Jiang, Leping Zhang, Xiaojun Huang, Michael Q. Zhang, Hong Wu

## Abstract

Recent studies have demonstrated that 3D genome alterations play important roles in tumorigenesis^1–3^, including the development of hematological malignancies^4–7^. However, how such alterations may provide key insights into T-lineage acute lymphoblastic leukemia (T-ALL) patients is largely unknown. Here, we report integrated analyses of 3D genome alterations and differentially expressed genes (DEGs) in 18 newly diagnosed T-ALL patients and 4 healthy T cell controls. We found that 3D genome organization at the compartment, topologically associated domains (TAD) and loop levels as well as the gene expression profiles could hierarchically classify different subtypes of T-ALL according to the T cell differentiation trajectory. Alterations in the 3D genome were associated with nearly 45% of the upregulated genes in T-ALL. We also identified 34 previously unrecognized translocations in the noncoding regions of the genome and 44 new loops formed between translocated chromosomes, including translocation-mediated enhancer hijacking of the *HOXA* cluster. Our analysis demonstrated that T-ALLs with *HOXA* cluster overexpression were heterogeneous clinical entities, and ectopic expressions of the *HOXA11*-A*13* genes, but not other genes in the *HOXA* cluster, were associated with immature phenotypes and poor outcomes. Our findings highlight the potentially important roles of 3D genome alterations in the etiology and prognosis of T-ALL.

---

T-ALL is an aggressive hematological malignancy caused by genetic and epigenetic alterations that affect normal T cell development^8^. Recent whole-exome and RNA sequencing analyses of large T-ALL cohorts are focused on the coding region of the genome and have identified novel driver mutations, dysregulated transcription factors and pathways in T-ALLs^9–12^. To determine whether alterations in the 3D genome organization are associated with malignant transformation of T-ALL, we conducted BL-Hi-C^13^ analysis using purified primary leukemic blasts from 18 newly diagnosed T-ALL patients, including 8 early T-cell precursor ALL (ETP ALL) and 10 non-ETP ALL, two clinical subtypes of T-ALL, as well as normal T cells from 4 healthy volunteers (Supplementary Fig. 1a). The maximum resolutions of the chromatin contact maps for ETP, non-ETP ALL and normal samples were approximately 3.5, 3.5 and 10 kb, respectively (Supplementary Table 1).

Principal component analysis (PCA) at the levels of the compartment, TAD and loop structures demonstrated that the T-ALL samples could be separated from the control samples by PC1, while ETP and non-ETP ALL could be separated by PC2 at all three architectural levels (Fig. 1a, upper panels) and be further delineated by hierarchical clustering analysis (Fig. 1a, lower panels). Detailed comparisons of the 3D chromosomal organizations of the T-ALL patients and the healthy controls revealed 1.59% of A-to-B and 1.38% of B-to-A T-ALL-associated switches at the compartment level (Supplementary Fig. 1b), 11% (377/3421) T-ALL-specific TAD boundaries (Supplementary Fig. 1c), and 6% (2330/38464) enhanced and 11% (4073/38464) reduced loops (Supplementary Fig. 1d), implying that the global 3D genome alterations could link to the etiology of T-ALL development.

**Fig. 1.**
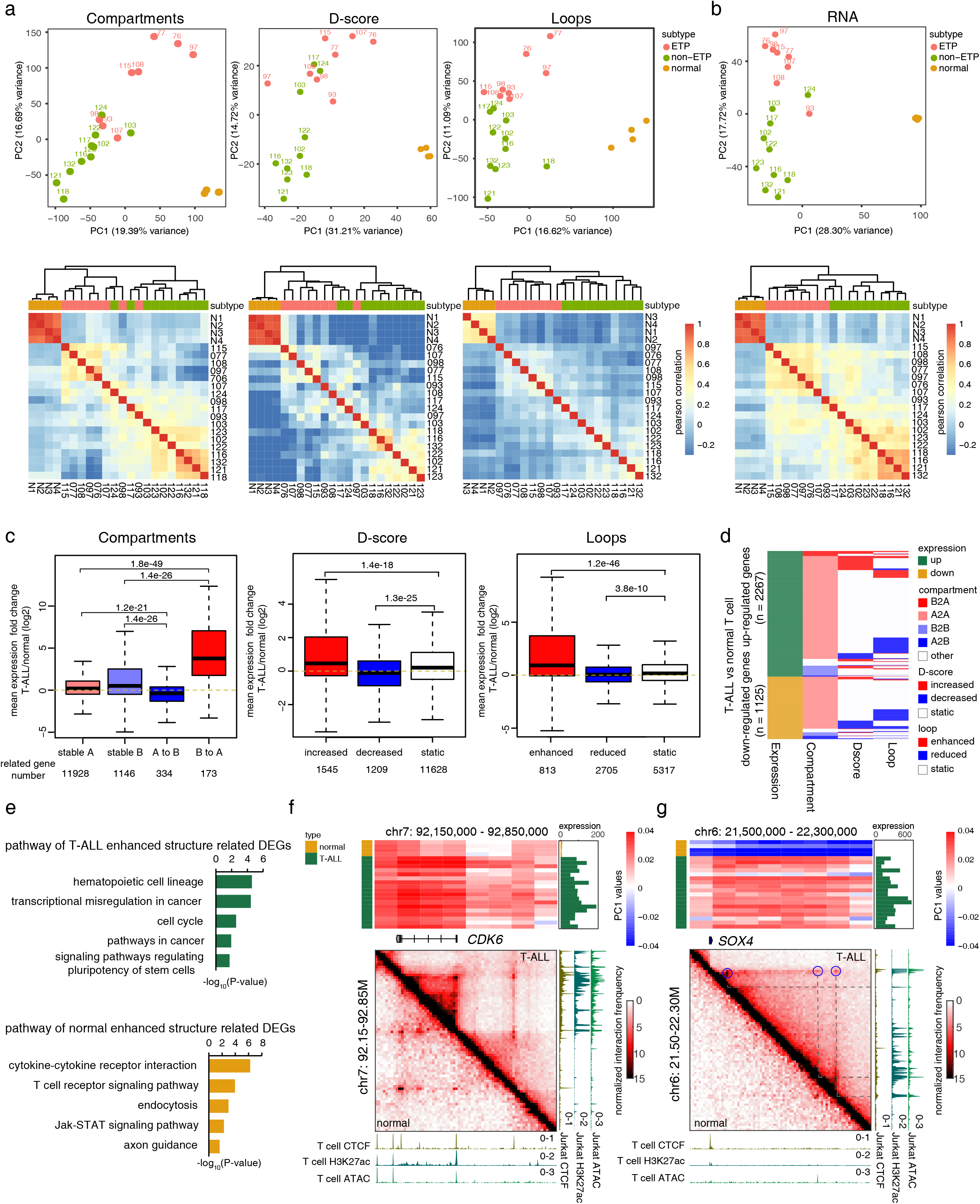
Global 3D genome architectures in T-ALLs. (a) PCA (upper) and unsupervised hierarchical clustering analysis (lower) of compartment, domain score (D-score) and loop in normal T cells and T-ALLs. (b) PCA (upper) and unsupervised hierarchical clustering analysis (lower) of gene expression profiles in normal T cells and T-ALLs. (c) Association of DEGs and genomic alterations at the levels of compartment, TAD and loop. Red, upregulation; blue, downregulated. The p values were calculated using Wilcoxon rank sum test. (d) A summary of DEGs and their corresponding chromatin structure changes. (e) KEGG analysis for enriched pathways in T-ALLs and normal T cells based on DEGs that are associated with chromatin structural changes. (f) and (g) Top, heatmaps show the compartment scores for each sample across the genomic regions of the *CDK6* and *SOX4* loci, respectively. Bar plots on the right show the gene expression level of *CDK6* and *SOX4* for each sample. Bottom, Hi-C contact maps of the *CDK6* and *SOX4* loci in normal T cell and T-ALL; blue circles: enhanced loops in T-ALL. ATAC-seq tracks and ChIP-seq tracks of CTCF and H3K27ac of normal T cells and T-ALL Jurket cells are also included.

RNA-seq analysis was also performed on each sample to investigate the impact of 3D genome alterations on gene expression in the T-ALLs. PCA and hierarchical clustering showed that the transcriptome changes were highly consistent with the corresponding 3D genome structural changes (comparing Fig. 1a and 1b) and hence could serve as a biological readout for the functional impact of 3D genome alterations in T-ALL development.

Approximately 29% (996/3392) of the DEGs were associated with 3D genome alterations (Supplementary Table 2). Among them, the genes associated with the B-to-A compartment switches, increased domain scores (D-score) and the corresponding enhanced loops were mostly upregulated (Fig. 1c, red bars and Fig. 1d), and were functionally enriched in pathways such as hematopoietic cell lineage, transcriptional misregulation in cancer and cell cycle (Fig. 1e). In contrast, the genes associated with the A-to-B compartment switches, decreased D-scores and reduced loops were mostly downregulated (Fig. 1c, blue bars and Fig. 1d), and these genes were enriched in pathways such as cytokine-cytokine receptor interaction and T cell receptor signaling (Fig. 1e).

Among the upregulated DEGs, *CDK6* is a potential target for T-ALL treatment^14^. The *CDK6* locus exhibited a strong intra-TAD interaction, and its expression was upregulated in all T-ALL samples (Fig. 1f; Supplementary Fig. 1e). Similarly, several upregulated oncogenic driver genes and T-ALL-associated transcription factors, such as *MYB*, *MYCN*, *BCL11A*, *SOX4* and *WT1*, also had increased D-scores (Supplementary Fig. 1f).

While the majority of the DEGs were associated with structural changes at 1 or 2 levels, transcription factor *SOX4*, *WT1* and *TFDP2* had structural alterations at all 3 levels (Fig. 1g, Supplementary Fig. 1g and Supplementary Table 2). Importantly, when RcisTarget^15^ was used to predict the key transcription factors that might control the upregulated genes in the T-ALLs, *SOX4*, *WT1* and *TFDP2* were among the top-ranking candidates that potentially regulated approximately 28% of the upregulated genes (Supplementary Table 3). After eliminating the overlapping upregulated DEGs, we estimated that 3D genome alterations could associate with approximately 45% of the upregulated genes in T-ALL, of which 56% could be directly related to structural alterations, while the rest could be due to structure alteration-mediated dysregulated transcription factors.

ETP ALL is a subclass of T-ALL stalled at the early T progenitor stage, while non-ETP ALLs are blocked at the later T cell differentiation stages^16^. Our 3D genome landscape analyses could separate the ETP ALL samples from the non-ETP ALL samples (Fig. 1a), suggesting that the chromosomal organizations of T-ALL may represent different “frozen stages” of T cell development^17^. To test this hypothesis, we projected the T-ALL samples onto the T cell developmental trajectory defined by RNA-seq analysis^18^. PCA revealed that most of the ETP ALL samples were arrested at the immature stage, corresponding to the LMPP to Thy1 stages, while the non-ETP samples were arrested at the Thy2 to Thy4 stages (Fig. 2a). Since the ETP and non-ETP ALLs can be better separated at the loop level (Fig. 1a), we further analysed the differences in loop structures between ETP and non-ETP ALL samples and identified 1820 enhanced and 831 reduced loops in ETP (Fig. 2b). When plotting gene expression changes between ETP and non-ETP ALL against the combined p-value of the loop strength and D-score changes, we found a strong positive correlation (Pearson’s correlation coefficient 0.685; Fig. 2c). Approximately 20% and 16% of the upregulated genes in ETP and non-ETP ALL, respectively, harbored chromatin structure changes, including key transcription factors or oncogenes, such as *CEBPA*, *MYCN* and *LYL1* for ETP and *LEF1*, *TCF12* and *PAX9* for non-ETP ALL (Fig. 2c and Supplementary Table 4).

**Fig. 2.**
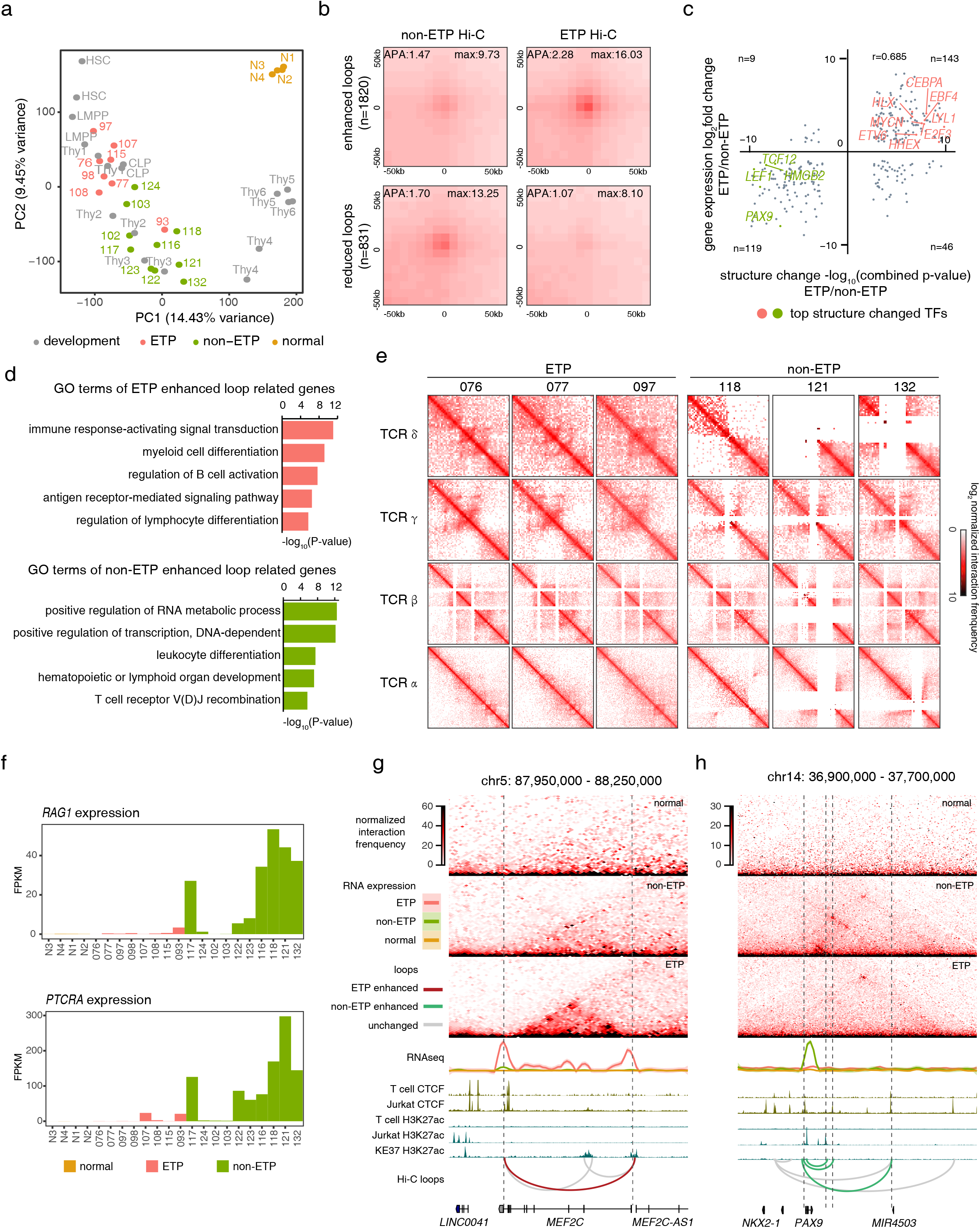
ETP and non-ETP ALLs have different loop structures. (a) PCA shows the association between ETP and non-ETP ALL gene expression profiles and T cell developmental trajectory. (b) APA plots for loops that are enhanced (top) or reduced (bottom) in ETP compared with non-ETP ALL. (c) Scatterplot shows the correlation between gene expression changes and structural changes. The structural change was defined by the combined p-value of D-score and loop strength change. Top ETP and non-ETP ALL-associated transcription factors with structural changes are highlighted in red and green, respectively. (d) GO terms for genes adjacent to enhanced loop anchors in ETP (top) and non-ETP (bottom). (e) Hi-C contact maps for the TCR genomic regions in ETP and non-ETP ALLs. (f) *RAG1* and *PTCRA* expression levels. g) and (h) Hi-C contact maps for TADs enclosing the genomic loci of ETP expressed *MEF2C* (g) and non-ETP expressed *PAX9* (h). Anchors of differential loops are labeled with dashed lines.

Gene ontology analysis further revealed that genes associated with the ETP ALL enhanced loops were enriched in immune response-activating signal transduction, myeloid cell differentiation and regulation of B cell activation, consistent with the definition of ETP ALL (Fig. 2d). Genes associated with the non-ETP ALL enhanced loops were enriched in terms such as positive regulation of RNA metabolism, transcription, and TCR V(D)J recombination (Fig. 2d). The lack of TCR rearrangement in most of the ETP ALL samples (Supplementary Fig. 2a) and different rearrangements in individual non-ETP samples (Supplementary Fig. 2b) could be easily observed from the Hi-C maps (Fig. 2e). Sample 093 was a unique case as it fell between ETP and non-ETP ALL (Fig. 1a and 1b; Fig. 2a) and had significant TCR rearrangement (Supplementary Fig. 2a). We also observed a lack of *RAG1* and *PTCRA* expression in most of the ETP ALL samples, which are essential for TCR V(D)J rearrangements (Fig. 2f).

The 3D genome alteration analysis also provided a potential explanation for ETP and non-ETP ALL-specific transcription factor expressions. For example, we detected subtype-specific loops and expression patterns in the *MEF2C* locus in ETP and the *PAX9* locus in the non-ETP ALL samples, respectively, which were associated with H3K27ac and CTCF marks in the ETP cell line KE37 and non-ETP cell line Jurkat (Fig. 2g and 2h).

Chromosomal translocation is one of the major driving forces for tumorigenesis^19^, especially for leukemia^8^. By adapting hic_breakfinder^20^, we identified 46 translocations in 14/18 T-ALL samples (Supplementary Table 5), of which 34 were novel events and 26 were interchromosomal events (Fig. 3a, red lines). Among 78 unique breakpoints identified, 47% were located in noncoding regions, and 66% were located in the stable A compartment (Supplementary Fig. 3a). These newly identified translocations not only influenced the expression of the nearest genes (Fig. 3a) but also resulted in the formation of 44 new loops across the translocated chromosomes, which we named translocation-mediated loops (Supplementary Fig. 3b; Supplementary Table 6). Interestingly, the ends of these translocation-mediated loops tend to anchor at the pre-existing loop anchors and CTCF binding sites (Fig. 3b). Importantly, nearly 78% of the translocation-mediated loops with CTCF motifs were linked to pairs of convergently orientated CTCF motifs (Fig. 3c), indicating that these loops may be mediated by loop extrusion mechanism^21–23^.

**Fig. 3.**
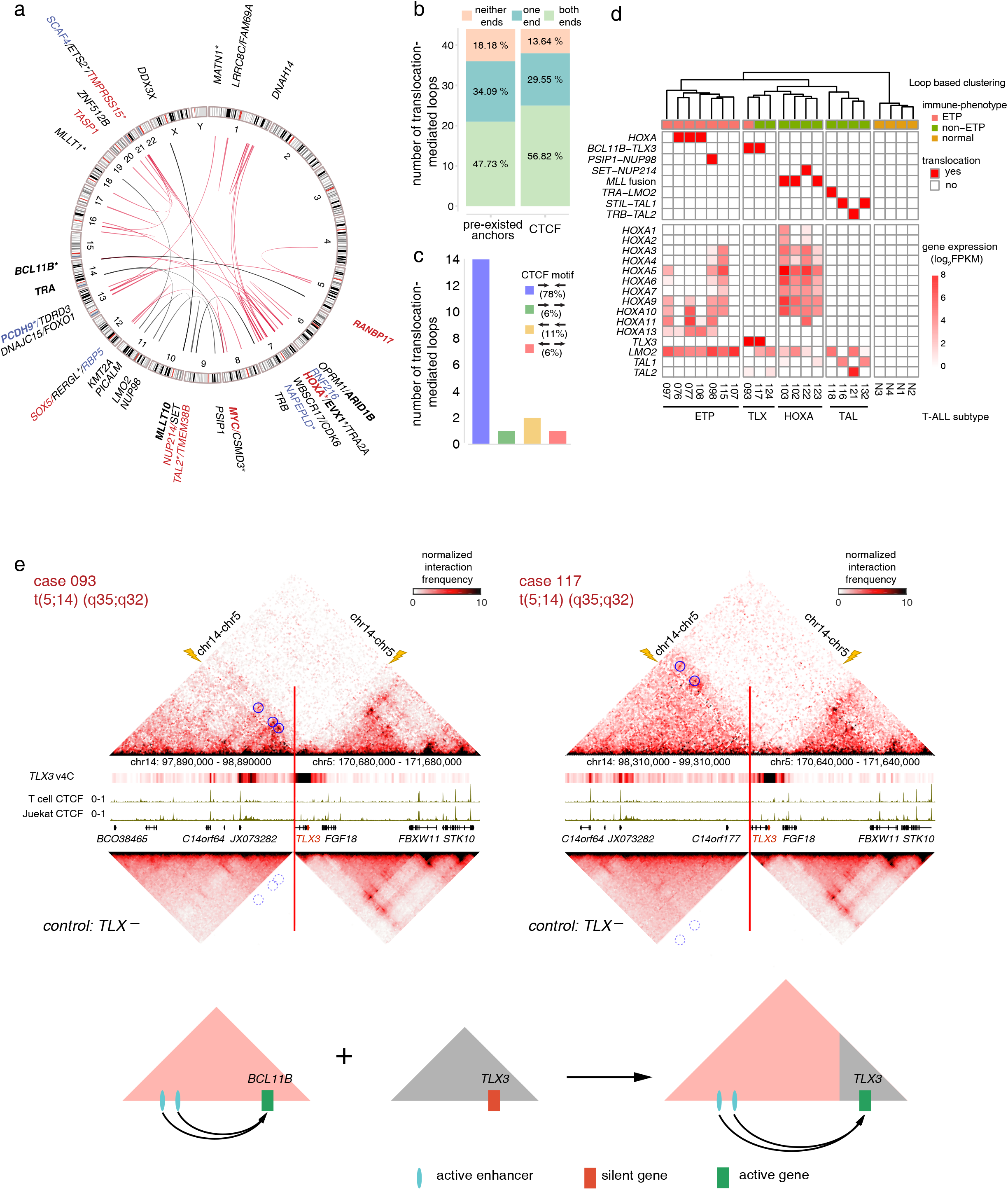
Chromosomal rearrangements in T-ALLs. (a) The genomic landscape of translocations discovered by Hi-C. The novel translocation partners are connected by red lines and known translocations are shown by black lines. Breakpoints nearest genes with increased or decreased expressions are highlighted by red and blue, respectively. Recurrent breakpoints nearest genes are labeled in bold. The genes closest to the breakpoint in noncoding regions are marked by star symbols. (b) Majority of the translocation-mediated loop anchors are pre-existing loop anchors and contains CTCF binding sites. (c) Majority of translocation-mediated loops have convergent CTCF motif orientation. (d) The loop-based clustering overlaps with leukemogenic transcription factor-or translocations-based clustering in T-ALLs. *STIL-TAL1* fusions were detected by RT-qPCR, the other translocations were detected by Hi-C. (e) Upper panels: Hi-C heatmaps for cases with *BCL11B-TLX3* translocation (top), visual 4C plots generated from Hi-C contact maps using *TLX3* promoter as the viewpoint (middle) and averaged Hi-C heatmaps for cases without *BCL11B-TLX3* translocation as controls (bottom). Breakpoints are marked by the red lines and yellow lightning bolts. Translocation-mediated loops are highlighted by blue circles, corresponding loop locations at controls are highlighted by dotted line circles. Lower panels: a schematic illustration for the consequence of the *BCL11B-TLX3* translocation.

We then investigated the potential mechanisms underlying translocation-mediated T-ALL classification and gene activation. Clinically, non-ETP ALL can be further classified into the HOXA, TLX and TAL subtypes according to their gene expression profiles^24^. Notably, there was a complete match between loop-based hierarchical clustering and T-ALL subtypes, which were signified by chromosomal translocation-mediated dysregulation of T-ALL-associated transcription factors (Fig. 3d).

Translocations can activate T-ALL-associated transcription factors via either “cis” or “trans” mechanisms. The “cis” mechanism involved translocation-mediated enhancer hijacking in the ETP, TLX and TAL subtypes (Fig. 3d and Supplementary Fig. 3b), of which the ectopically expressed genes, such as *TLX3*, hijacked the enhancers from the translocated *BCL11B* gene via translocation-mediated loops (Fig. 3e and Supplementary Fig. 3c-d). The “cis” mechanism included *BCL11B-TLX3*, *TRB-TAL2*, and 3 novel *HOXA* translocations identified in this study (Supplementary Fig. 3b and Supplementary Table 6; see Fig. 5). Interestingly, most of the hijacked enhancers are from genes that are normally expressed during T cell development, such as *BCL11B* and *TRB*, which lead to ectopic expression of T-ALL-associated transcription factors in the T lineage and block normal differentiation (Supplementary Fig. 3d). The “trans” mechanism involves translocation-mediated gene fusions, such as the *PSIP1-NUP98*, *SET-NUP214*, and *MLL* (*KMT2A-MLLT1*, *PICALM-MLLT10*, and *DDX3X-MLLT10*) gene fusion events, which could epigenetically activate *HOXA* cluster gene expressions^25–28^. These results suggest that translocation-mediated enhancer hijacking or fusion events may drive ectopic transcription factor activation, leading to specific pathogenic gene expression profiles of the T-ALL subgroups.

The dysregulated *HOXA* cluster, which contains 11 genes, is a common feature of T-ALL^24^. However, whether *HOXA*-positive T-ALLs represent a homogeneous clinical entity has not been systematically studied. We conducted unsupervised hierarchical clustering based on *HOXA* gene expressions, which separated 15 T-ALL samples without *HOXA* translocation into *HOXA*-negative (HOXA^−^) and *HOXA*-positive expression (HOXA^+^) groups. Translocation-mediated fusion events could be detected in 5/7 HOXA^+^ samples. The HOXA^+^ T-ALLs could be further separated into 2 subgroups: the 3’HOXA^+^ or 5’HOXA^+^ subgroups, with respect to the location of the *HOXA* genes within the *HOXA* cluster (Fig. 4a). The expression patterns of the three *HOXA* translocation cases (HOXA-T, breakpoints shown in Fig. 4b) were closer to those in the 5’HOXA^+^ subgroup, characterized by ectopic *HOXA11-A13* expressions (Fig. 4a). Interestingly, the 5’HOXA^+^ and HOXA-T cases in our Hi-C study are associated with double negative (DN) and ETP phenotypes (Fig. 4a), suggesting that T-ALLs with ectopic *HOXA* cluster gene expression are heterogeneous entities.

**Fig. 4.**
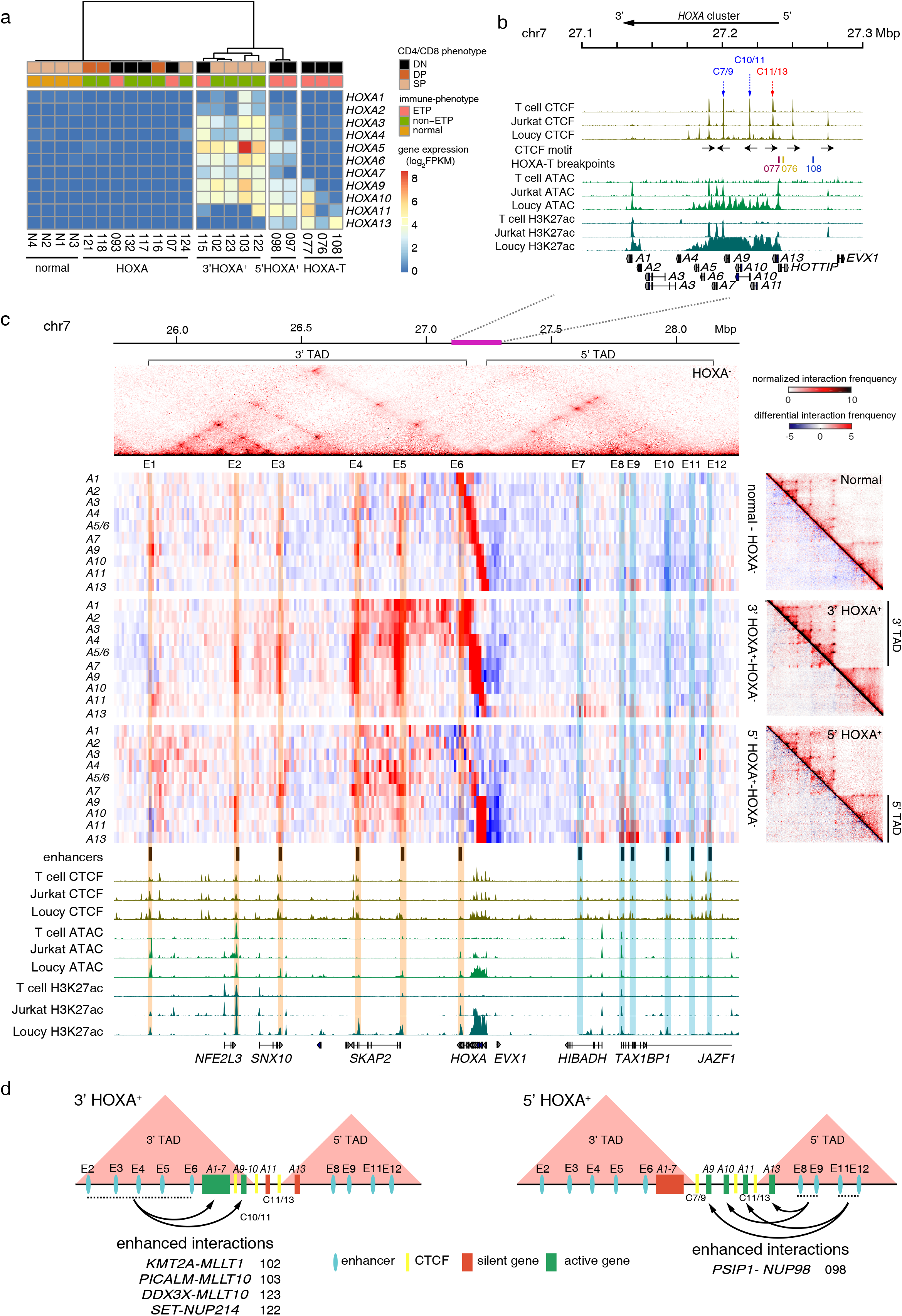
Chromatin interaction profile and expression patterns of the HOXA cluster in T-ALLs. (a) T-ALL subtypes with different *HOXA* gene expression patterns. HOXA-translocation (HOXA-T) negative cases are grouped by unsupervised hierarchical clustering based on the *HOXA* gene expressions. (b) ChIP-seq tracks of CTCF and H3K27ac and ATAC-seq tracks in T cell, HOXA^−^ Jurkat and HOXA^+^ Loucy T-ALL cells in the genomic region corresponding to the pink bar of Fig. 4c. CTCF motif orientations, 5’TAD and 3’TAD boundaries and HOXA-T breakpoints are also included. (c) Top: a Hi-C heatmap shows the average interaction intensity of HOXA^−^ cases in chr7: 25,750,000-28,250,000 (hg19), which includes *HOXA* gene cluster and its 3’ and 5’ TADs. Middle left: from top to bottom, Hi-C heatmaps show differential interaction intensities between normal T cell vs. HOXA^−^, 3’ HOXA^+^ vs. HOXA^−^ and 5’ HOXA^+^ vs. HOXA^−^ cases using visual 4C plot. Each line represents interactome of a *HOXA* gene shown on the left. *HOXA5* and *HOXA6* share one line as they located in the same bin. The main enhancers are highlighted with orange in the 3’ TAD and blue in the 5’ TAD. Middle right: Top right of each heatmap represents the averaged interaction intensities of normal T cell, 3’ HOXA^+^ cases and 5’ HOXA^+^ cases, respectively; bottom left of each heatmap represents the differential interaction intensities between each group and HOXA^−^ cases. Bottom: ChIP-seq tracks for CTCF and H3K27ac and ATAC-seq tracks in T cell, HOXA-Jurkat and HOXA^+^ Loucy T-ALL cells. (d) Schematic illustrations of the associations among 3D genomic interactions, *HOXA* cluster expressions and associated fusion events in 3’ HOXA cases (left) and 5’ HOXA cases (right).

The *HOXA* genes are transcriptionally repressed in normal T cells but can be transactivated in T-ALLs by fusion proteins that recruit histone methyltransferase DOT1L to the *HOXA* locus^25,29^. Although this mechanism uncovered how the *HOXA* cluster is activated, it cannot explain the diverse *HOXA* expression patterns associated with different fusion proteins. By integrating Hi-C maps with *HOXA* gene expression patterns, we found that the differential *HOXA* gene expressions were associated with different 3D genome organizations in samples without *HOXA* translocation. Hi-C maps and CTCF motif orientations showed that the 11 *HOXA* genes were partitioned between 2 TADs (Fig. 4b and Supplementary Fig. 4a): the CTCF binding site C11/13 was used as the 3’ boundary of the 5’ TAD in all samples, while the 5’ boundaries of the 3’ TADs varied among different samples: C7/9 was used by most of the HOXA^−^ (6/8) and all 5’HOXA^+^ samples (2/2), while C10/11 was used by most of the 3’HOXA^+^ samples (4/5) (red and blue arrows/lines, respectively; Fig. 4b and Supplementary Fig. 4a).

We further identified 6 enhancer regions in each TAD, labeled E1 to E12, which could interact with the *HOXA* cluster (Fig. 4c). Using HOXA^−^ cases as common denominators (Fig. 4c, top panel), we calculated the overall differential interaction intensities. Although there was no significant difference between the healthy controls and the HOXA^−^ cases in the 12 interaction regions, we found significantly enhanced interactions between E2-E6 and genes in the 3’HOXA subgroup, as well as between E8, 9, 11, 12 and genes in the 5’HOXA subgroups, either as a group average (Fig. 4c) or individually (Supplementary Fig. 4b-c). ChIP-seq analysis of the HOXA^+^ Loucy cell line indicated that these increased interactions may be correlated with gains in the H3K27ac histone mark (Fig. 4c). The KMT2A-MLLT1, PICALM-MLLT10, DDX3X-MLLT10 and SET-NUP214 fusions were associated with interaction dynamics and HOXA1-A10 expression in the 3’HOXA subgroup, while the PSIP1-NUP98 fusion event was associated with the 5’HOXA subgroup (Fig. 4d). These results suggest that different translocation-mediated fusion events may epigenetically and preferentially alter the 3D genome interactome within the *HOXA* cluster and control a specific set of *HOXA* gene expression.

To explain the gene expression patterns seen in the three HOXA-T samples, we mapped the breakpoints and found that all the breakpoints were located within the 5’ TAD, upstream of the *HOXA13* gene. The translocation partner breakpoints lie in the gene bodies of the *BCL11B* and *CDK6* genes on chromosome 14 and chromosome 7, respectively, as well as upstream of the *ERG* gene on chromosome 21 (Fig. 5 and Supplementary Fig. 5). By examining the TAD structures associated with the translocations, using HOXA^−^ samples as controls, we found that these translocations mediated new loop formations between the 5’ of the *HOXA* cluster and active enhancers associated with *BCL11B* and *ERG* genes, leading to ectopic expression of *HOXA13* in the case of 076 and *HOXA9-A13* in the case of 077 (blue circles for new loops and green bar graphs for gene expressions; Fig. 5a and b; Supplementary Fig. 3b). For the case of 108, the inter-TAD inversion led to the adoption of active CDK6 enhancers and new loop formation (Fig. 5c), causing ectopic expression of *HOXA11-A13* (Fig. 5c). These results suggest that translocations and inversion associated with the 5’ TAD of the *HOXA* cluster lead to dysregulated 5’ *HOXA* gene expressions through new loop formation-mediated enhancer hijacking. Again, the existence of the CTCF binding sites C11/13 in the case of 076, C7/9 in the case of 077, and C10/11 in the case of 108 (Fig. 5d) could insulate genes located in the 3’ TAD from the influence of the hijacked enhancers, leading to 5’HOXA-specific expression patterns (Fig. 4b).

**Fig. 5.**
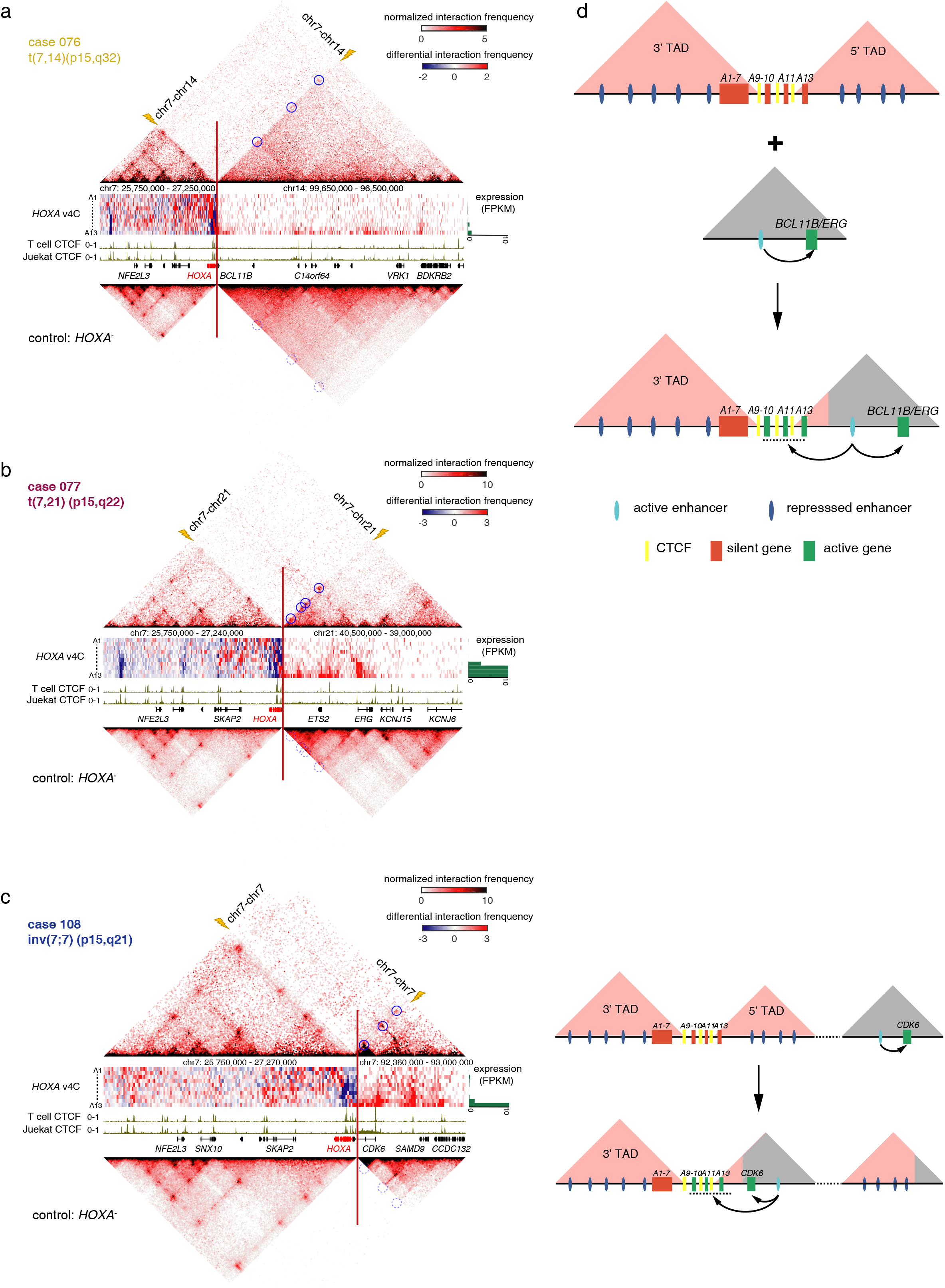
Translocation-mediated enhancer hijacking and ectopic *HOXA* gene expressions in T-ALLs. (a-c) Individual Hi-C heatmaps for three HOXA-T cases (top) and averaged Hi-C heatmaps of HOXA^−^ control cases (bottom). Middle panels are differential visual 4C plots between HOXA-T cases and HOXA^−^ using *HOXA1-A13* genes (from top to bottom) as the viewpoint. The green bar graphs on the right show the expression level of each *HOXA* gene. Breakpoints are marked by the red lines and yellow lightning bolts. Translocation-mediated loops are highlighted by blue circles, corresponding locations in control samples are marked by dotted circles. (d) Schematic illustrations of the consequences of *HOXA*-related translocations (upper) and inversion (lower).

We next investigated whether ectopic *HOXA* cluster expression was associated with poor prognosis. For this, we analyzed a cohort of T-ALL patients with outcome information (see our companion paper). We found that *HOXA11* or *HOXA13* positivity, alone or in combination, but not the expression of other *HOXA* genes, such as the previously used biomarker *HOXA9*, was associated with poor overall and event-free survivals in young adult and pediatric T-ALLs. In contrast, T-ALLs with *TAL1/2* positivity, which was mutually exclusive from that with *HOXA* positivity, were associated with better overall survival, while TLX positivity had no association with the outcome (Fig. 6a and Supplementary Fig. 6a-b). Compared to the *HOXA1-A10*^+^, *TAL*^+^ and *TLX*^+^ samples, the *HOXA11-A13*^+^ cases were characterized by DN and ETP phenotypes (Fig. 6b and Supplementary Fig. 6c), as well as a higher rate of JAK-STAT pathway gene mutations^9,30^ (Fig. 6c). In multivariate analysis, *HOXA13*^+^ status could serve as an independent predictor for the overall survival of pediatric and young adult T-ALLs (Fig. 6d).

**Fig. 6.**
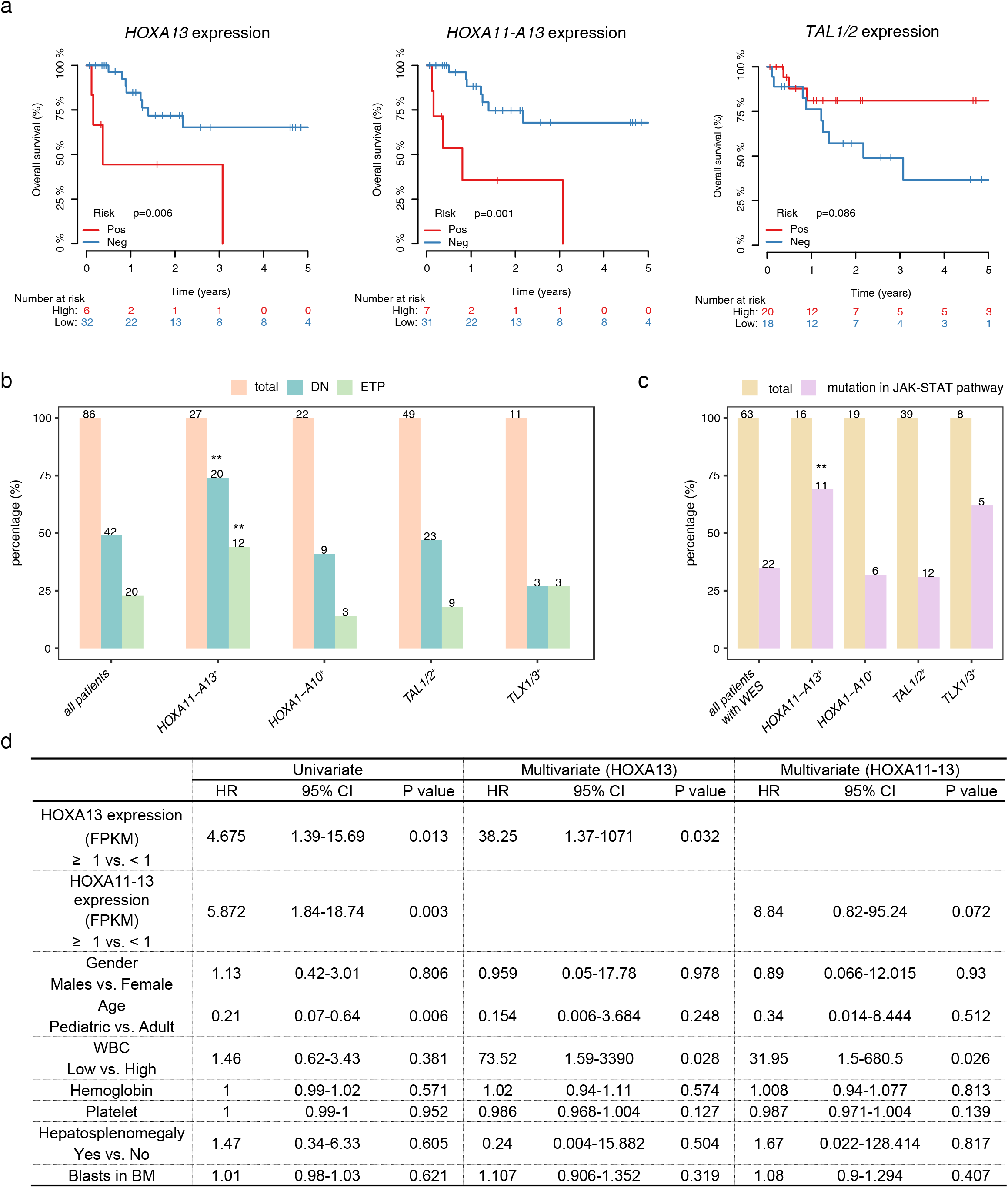
Ectopic *HOXA11*-*A13* expressions are correlated with poor outcomes in pediatric and young adult T-ALLs. (a) Kaplan-Meier overall survival curves for patients with (red) and without (blue) ectopic *HOXA13* expression (left), *HOXA11-A13* expressions (middle) and *TAL1/TAL2* expressions (right) in pediatric and young adult patients. (b) The associations between ectopic *HOXA11-A13* expression and DN and ETP phenotypes. *P*-values are calculated by Fisher exact test (**: p<0.01). (c) The proportion of cases with JAK-STAT pathway mutations in each T-ALL subgroup, *P*-values are calculated by Fisher exact test (**: p<0.01). (d) Univariable and multivariable analysis of overall survival. HR, hazard ratio; CI, confidence interval.

Taken together, our results imply that 3D genome alterations, in addition to previously identified oncogenic driver mutations and dysregulated pathways, may play critical roles in rewiring T-ALL transcriptome and regulating T-ALL development. The newly identified translocations and enhancer hijacking mechanism may provide a potential explanation for how T-ALL-associated transcription factors are ectopically activated and block differentiation at certain T cell developmental stages. Finally, we identified the associations of ectopic *HOXA11-A13* expression and poor survival of T-ALL, corresponding to the subtype currently incurable by standard treatments, and suggested that anti-JAK-STAT inhibitors may benefit this group of patients.

## Author contributions

LY, HZ and HW conceived the project; YC designed the Hi-C and RNA-seq experiments; LY, HZ, MS and WW performed the Hi-C and RNA-seq experiments; FC designed the bioinformatic pipelines and performed the Hi-C and RNA-seq integrated analyses, while BD conducted the survival analysis. QJ, LZ and XH contributed the clinical samples and data. LY, FC and BD generated the figures and tables. LY, FC and HW wrote the manuscript with help from all authors. XH was in charge of the clinical study; MZ and YC oversaw the bioinformatics analyses, and HW supervised the entire project.

## Acknowledgement

We thank Drs. Meng Lv, Yingjun Chang, and Yan Chang for sample collection; Dr. Cheng Li of Peking University for critically reviewing the manuscript. We also thank Yan Liu, Fei Wang and Xuefang Zhang from the National Center for Protein Sciences Beijing at Peking and Tsinghua Universities for assistance with FACS. This project was supported by the Peking-Tsinghua Center for Life Sciences, Beijing Advanced Innovation Center for Genomics at Peking University for HW and the National Natural Science Foundation of China (81602254 for LY, 31871343 for YC, 31671384 and 81890994 for YC and MZ). WW was supported by the Postdoctoral Fellowship of Peking-Tsinghua Center for Life Sciences.

## Materials and Methods

### Patients and samples

Eighteen T-ALL patient samples were collected under the protocol approved by the ethics committee of the Institute of Hematology at Peking University. The patient characteristics are described in Supplementary Table 7. ETP status was defined as previously published^16^. Leukemia blast cells were prepared by density-gradient centrifugation of bone marrow samples, and CD19^−^CD14^−^CD235^−^CD45^+^CD7^+^ cells were further purified by FACS analysis using anti-human antibodies for RNA-seq and Hi-C library preparations. Peripheral blood samples were obtained from four healthy donors under the approval of the ethics committee of Peking University. T cells were purified using the EasySep™ Direct Human T Cell Isolation Kit (StemCell Technology #19661).

### RNA-seq library preparation, data processing, and differential gene expression analysis

RNA-seq libraries were prepared with TruSeq RNA Library Prep Kit v2 (Illumina). Paired-end RNA-seq reads of the 18 patients and 4 healthy controls were generated with an average depth of 15 million read pairs. Reads were aligned to the hg19 genome with TopHat (v2.1.0) using default settings^31^. Duplicates were removed, and aligned reads were calculated for each protein-coding gene using HTSeq^32^, followed by FPKM transformation by normalizing gene exon length and sequencing depth. Raw RNA-seq data for Loucy and Jurkat were downloaded from the GEO database and analyzed as described above.

DESeq2^33^ was applied to identify the differentially expressed genes with FDR <0.01 and fold change >2. Genes with fewer than 5 reads in 20% of the samples or with mean reads fewer than 2 were excluded. Signal tracks were generated by using BEDTools^34^ genomeCoveragebed to produce bedGraph files scaled to 1 million reads per data set. Then, the UCSC Genome Browser utility^35^ bedGraphToBigWig was used with default parameters to generate bigwig files.

### ChIP-seq, ATAC-seq data processing and motif analysis

ChIP-seq reads were mapped to the hg19 genome with Bowtie2^36^ (v2.3.5) using default parameters, while ATAC-seq reads were mapped with Bowtie2 using parameter -X 2000 --no-mixed --no-discordant --no-unal. Aligned reads were filtered for a minimum MAPQ of 20, and duplicates were removed using SAMtools^37^. Signal tracks and peaks were generated by using the -SPMR option in MACS2^38^. Then, the UCSC Genome Browser utility bedGraphToBigWig was used with default parameters to transform the bedgraph files to bigwig files. FIMO^39^ was used to detect the 20 bp CTCF motif from the Homer motif database in Loucy and Jurkat CTCF peaks with default parameters.

### Hi-C and Hi-C data processing

Hi-C was performed on one million cells/sample, according to the BL-Hi-C protocol^13^. Raw BL-Hi-C reads were processed by the in-house HiCpipe framework, which integrated several Hi-C analysis methods for to generate multiple features of the Hi-C data. In particular, ChIA-PET2^40^ was used to trim the bridge linkers and HiC-Pro^41^ to align reads, filter artifact fragments, and remove duplicates; Juicer^42^ was applied to the resulting uniquely mapped contacts to generate individual or merged Hi-C files that could be deposited as contact matrices with multiple resolutions. Knight-Ruiz^43^ (KR)-normalized matrices were used in the compartment and TAD analyses.

### Compartment and TAD analysis

The compartment was calculated with the eigenvector command of Juicer under 100 kb resolution KR normalized Hi-C matrices. For every 100 kb bin, A or B compartments were defined by the over 70% sample majority rule.

TAD boundaries were calculated by the Insulation score method^44^ (with parameters: -is 1000000 -ids 200000 -im mean -nt 0.1) on pooled 40 kb Hi-C matrices of the healthy T cell controls, ETP and non-ETP samples. The resulting TAD boundaries were merged and assigned with relative insulation scores of all samples calculated from HiCDB^45^. Differential TAD boundaries were defined with a t-test FDR<0.01 and a difference between cases and controls higher than 50% quantile of the overall difference.

A TAD was defined when its boundaries were detected in at least two conditions among normal T cells, ETPs and non-ETPs. The domain score^46^ was calculated in each sample by dividing the intra-TAD interactions with all interactions connected to the corresponding TAD. Differential domain scores were calculated with a t-test FDR<0.01 and fold change higher than 70% quantile of the overall fold change.

### Loop detection and differential loop calling

Loops were called by HiCCUPS^42^ at 5 kb and 10 kb resolutions with default parameters (except -d 15000,20000) for pooled Hi-C matrices of the healthy T cell controls, ETP and non-ETP samples, respectively. The differential loop detection method was adapted from Douglas *et al*^47^. Loops were split into two distance ranges (> or < 150 kb) to minimize potential bias (Rubin *et al*.^48^). Differential loops were called within each range (FDR < 0.1) and then combined.

### Loop aggregation and functional analysis

Aggregate peak analysis (APA) plots were generated to assess the quality of loop detection and explore the characteristics of different loop classes by the Juicer APA command^42^ under 5 kb resolution. Its output matrix was normalized by the loop number that contributed to the matrix generation. For analysis of the function of dynamic loops between non-ETP and ETP, the loop anchors were analyzed by GREAT^49^ (v3.0.0) using the nearest gene within 100 kb to generate the enriched biological process. In other sections, genes related to loops were determined if their promoter (5 kb around TSS) overlapped the loop anchors, and DAVID^50^ 6.8 was used for KEGG pathway enrichment analysis.

### Visualization and V4C plot generation

Tracks of Hi-C maps and ChIP-seq were generated by pyGenomeTracks^51^. Hi-C maps of each condition were normalized by its cis interaction pairs. A visual 4C (V4C) plot for specific loci was generated as the interaction s related to the corresponding viewpoint under 10 kb resolution.

### Translocation and translocation-mediated loop detection with Hi-C

hic_breakfinder^20^ was adapted to detect translocations in 18 T-ALL patient samples. After we filtered the “translocations” also detected in normal controls, the remaining translocations were manually assessed, and the precise breakpoints were determined. As the average depth of patient Hi-C samples is 486 million read pairs and the read length is 150 bp, Hi-C raw data were treated as single ends to refine the breakpoint locations to single base-pair resolution. Any single ends that could be mapped to two different chromatins without the BL-Hi-C bridge linker in between were chimeric reads. The chimeric reads detected from the BL-Hi-C data overlapped with the aforementioned translocations at 5~20 kb resolution. For each translocation, the exact locations (single base-pair resolution) supported by more than 3 chimeric reads were identified as the actual breakpoints and are reported in Supplementary Table 5.

As translocation-mediated loops were hard to identify by loop detection tools designed for intrachromosomal loop detection and easy to capture by visualization, their locations were manually recorded on interchromosome Hi-C maps with the help of Juicebox^52^, which is an interactive visualization software.

### Translocation, translocation-mediated loop annotation and visualization

The nearest genes to translocation breakpoints were determined by BEDTools. Known translocations were collected from references 2 and 4 and ChimerPub^53^. A translocation was considered novel if any of the breakpoints was not near any known breakpoint within a 100 kb distance. For translocation-mediated loops, the genes with a promoter or gene body overlapping the loop anchors were annotated as the associated genes. A gene near a breakpoint was considered upregulated if its FPKM was >1- and 2-fold higher than the control samples without nearby breakpoints. Hi-C heatmaps and Visual 4C plots of the reassembled chromatin were generated by MATLAB code.

### Statistics

Specific statistical analyses are described in each section. In general, the Wilcoxon rank sum test was employed in R for comparisons of distributions. Survival analysis was performed by a Cox regression model using overall and event-free survival as outcomes. Overall survival was defined as the time from diagnosis to death from any cause. Event-free survival was defined as the time from diagnosis to treatment failure, relapse, or death from any cause. The proportional hazard assumption was tested. Variables tested in the multivariable Cox regression model were sex, age (pediatric vs. adult), white blood cell counts, hemoglobin levels, platelet counts, hepatosplenomegaly, percentage of blasts in the bone marrow and MRD status. Eighty-six patient samples with RNA-seq data were used for DN- and ETP- enrichment analyses, of which 38 samples under the age of 40 with outcome and 63 samples with whole exon sequencing (WES) data were used for survival and mutation analyses, respectively.

### Code availability

Custom scripts described in the Online Methods will be made available upon request.

### Data availability

All sequencing data are available through the Gene Expression Omnibus (GEO) via accession GSE. Accession codes of the published data used in this study are as follows: CTCF ChIP-seq of CD4+ T cell and Jurkat cell line, GSE12889; CTCF ChIP-seq of Loucy cell line, GSE123214; ATAC-seq of CD4+ T cell, GSE87254; ATAC-seq of Jurkat cell line, GSE115438; H3K27ac ChIP-seq of CD4+ T cell, GSE122826; H3K27ac ChIP-seq of Jurkat cell line, GSE68978; H3K27ac ChIP-seq of Loucy cell line, GSE74311; RNA-Seq of Loucy cell line, GSE100694; RNA-seq of T cell development, GSE69239. The raw sequence data reported in this paper have been deposited in the Genome Sequence Archive^54^ in BIG Data Center^55^, Beijing Institute of Genomics (BIG), Chinese Academy of Sciences, under accession number HRA000113 that is publicly accessible at https://bigd.big.ac.cn/gsa.

