## Supplementary Figure 1 for "3D Genome Analysis Identifies Enhancer Hijacking Mechanism for High-Risk Factors in Human T-Lineage Acute Lymphoblastic Leukemia"

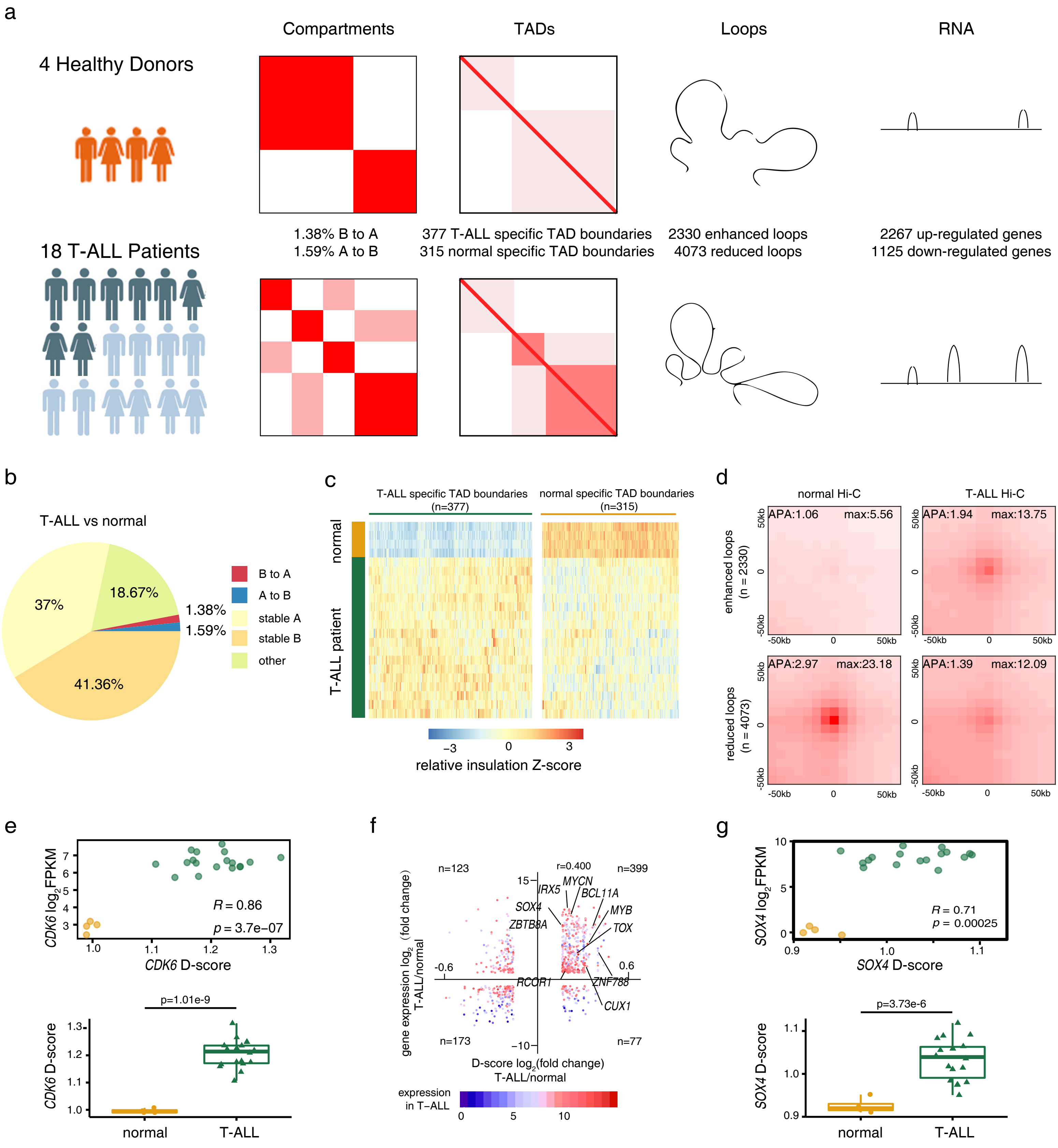
a
Compartments
TADs
Loops
RNA
4 Healthy Donors
18 T-ALL Patients
1.38% B to A
1.59% A to B
377 T-ALL specific TAD boundaries
315 normal specific TAD boundaries
2330 enhanced loops
4073 reduced loops
2267 up-regulated genes
1125 down-regulated genes
b
T-ALL vs normal
37%
18.67%
1.38%
1.59%
41.36%
B to A
A to B
stable A
stable B
other
c
T-ALL specific TAD boundaries (n=377)
normal specific TAD boundaries (n=315)
normal
T-ALL patient
−3 0 3
relative insulation Z-score
d
normal Hi-C
T-ALL Hi-C
enhanced loops (n = 2330)
reduced loops (n = 4073)
APA:1.06 max:5.56
APA:1.94 max:13.75
APA:2.97 max:23.18
APA:1.39 max:12.09
-50kb 0 50kb
e
CDK6 log2FPKM
CDK6 D-score
R = 0.86
p = 3.7e−07
p=1.01e-9
normal
T-ALL
f
n=123
r=0.400
n=399
n=173
n=77
IRX5
MYCN
BCL11A
SOX4
MYB
ZBTB8A
TOX
RCOR1
ZNF788
CUX1
-0.6
0.6
15
−10
gene expression log2 (fold change) T-ALL/normal
D-score log2(fold change) T-ALL/normal
expression in T−ALL
0 5 10
g
SOX4 log2FPKM
SOX4 D-score
R = 0.71
p = 0.00025
p=3.73e-6
normal
T-ALL

**Supplementary Fig. 1 Global 3D genome architecture in T-ALLs.** (a) Schematic illustrations of the study design and key findings comparing normal T cell and T-ALL. 22 samples, including 4 healthy donors (orange) and 18 T-ALL patients (8 ETP ALL, dark blue; 10 non-ETP ALL, light blue), are used for integrated genomic and transcriptome analyses. (b) Distributions of genomic regions associated with various compartment changes. (c) Heatmaps showing the relative insulation score of T-ALL specific TAD boundaries (left) and normal specific TAD boundaries (right). (d) APA plots for loops that are enhanced (up) or reduced (down) in T-ALLs compared with normal controls. (e) and (g) Upper: domain scores are plotted against gene expression of *CDK6* (e) and *SOX4* (g) genes. Lower: Quantification of the domain scores across TAD region encompassing the *CDK6* and *SOX4* genes in normal T cells and T-ALL samples. Statistical significance is calculated using t-test. (f) Domain score changes are plotted against gene expression changes in all DEGs between T-ALLs and normal T cell controls.
