## Supplementary Figure 2 for "3D Genome Analysis Identifies Enhancer Hijacking Mechanism for High-Risk Factors in Human T-Lineage Acute Lymphoblastic Leukemia"

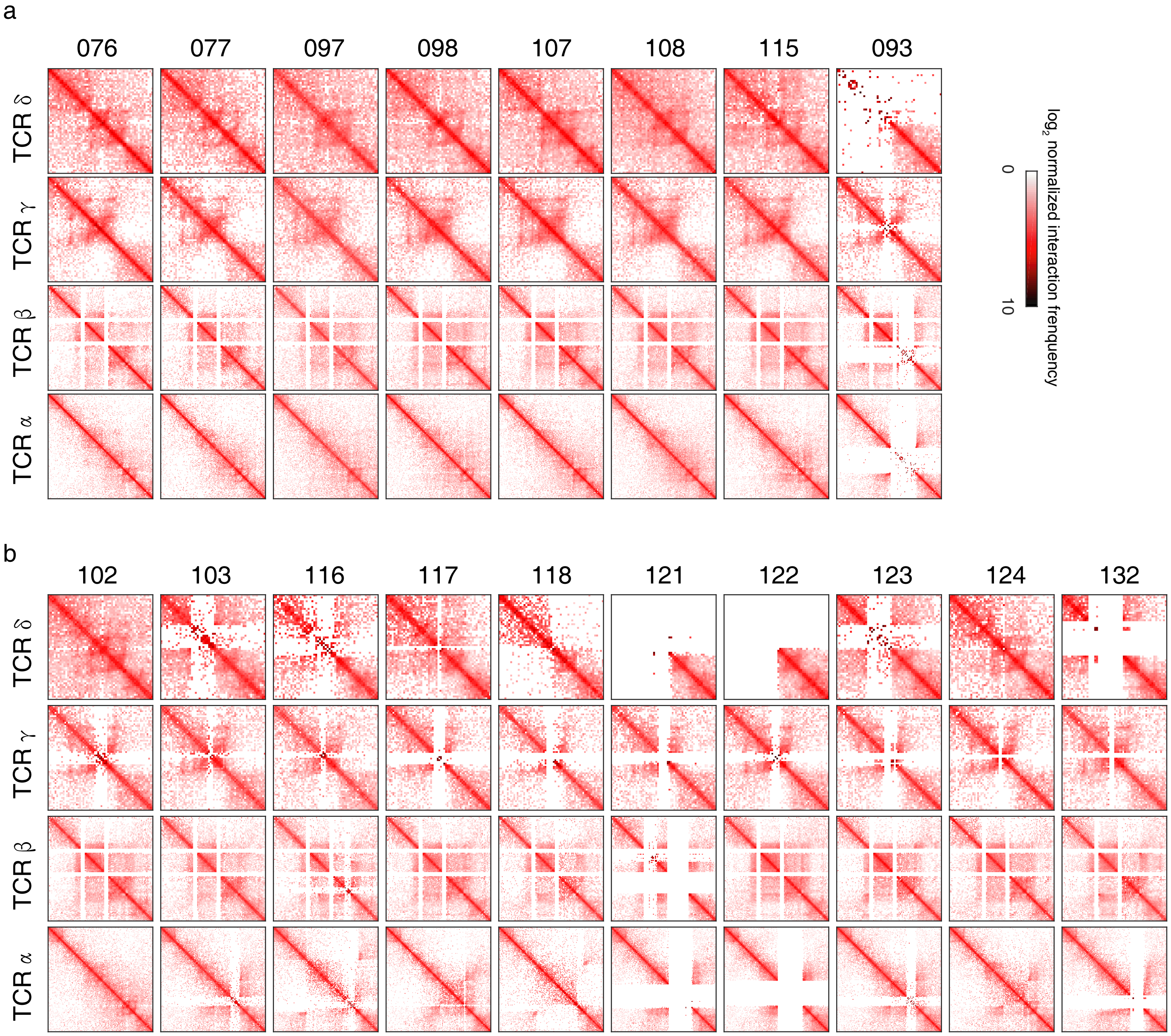


**Supplementary Fig. 2 ETP and non-ETP ALLs have different loop structures.** Hi-C contact maps for various TCR regions in 8 ETP (a) and 10 non-ETP cases (b).
