## Supplementary Figure 3 for "3D Genome Analysis Identifies Enhancer Hijacking Mechanism for High-Risk Factors in Human T-Lineage Acute Lymphoblastic Leukemia"

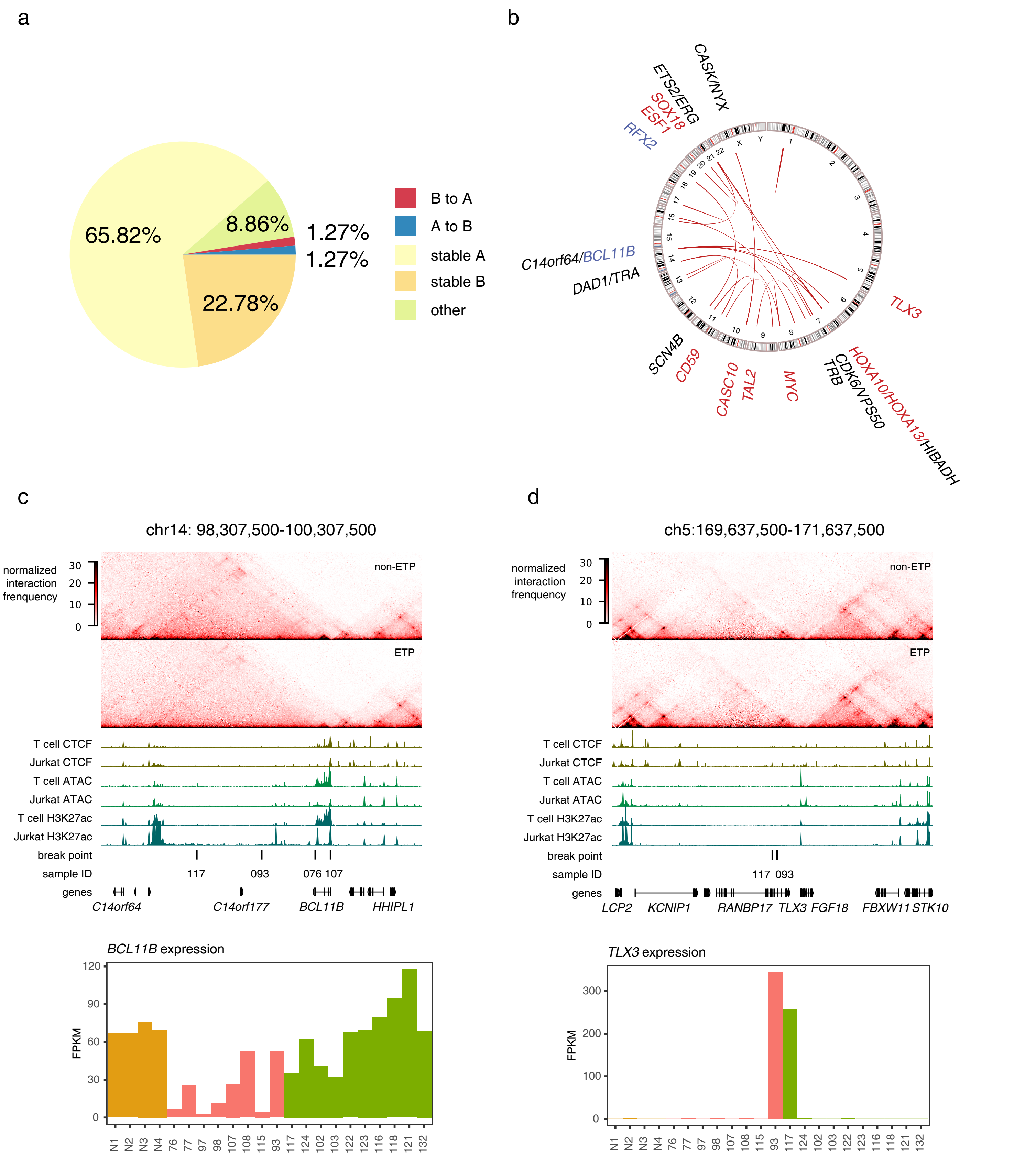
a
65.82%
8.86%
1.27%
1.27%
22.78%
B to A
A to B
stable A
stable B
other
b
CASK/NYX
ETS2/ERG
SOX18
ESF1
RFX2
C14orf64/BCL11B
DAD1/TRA
TLX3
SCN4B
CD59
CASC10
TAL2
MYC
TRB
CDK6/VPS50
HOXA10/HOXA13/HIBADH
c
chr14: 98,307,500-100,307,500
normalized interaction frenquency
non-ETP
ETP
T cell CTCF
Jurkat CTCF
T cell ATAC
Jurkat ATAC
T cell H3K27ac
Jurkat H3K27ac
break point
sample ID
117
093
076 107
genes
C14orf64
C14orf177
BCL11B
HHIPL1
BCL11B expression
FPKM
d
ch5:169,637,500-171,637,500
normalized interaction frenquency
non-ETP
ETP
T cell CTCF
Jurkat CTCF
T cell ATAC
Jurkat ATAC
T cell H3K27ac
Jurkat H3K27ac
break point
sample ID
117 093
genes
LCP2
KCNIP1
RANBP17
TLX3
FGF18
FBXW11
STK10
TLX3 expression
FPKM

**Supplementary Fig. 3 Chromosomal rearrangements in T-ALLs.** (a) Distributions of breakpoint locations in various compartments. (b) Novel translocation-mediated loops discovered by Hi-C. Genes adjacent to loop anchors with increased and decreased expressions are marked by red and blue, respectively. (c) and (d) Upper: Hi-C contact maps for TADs enclosing the genomic loci of *BCL11B* (c) and *TLX3* (d) in non-ETP and ETP samples. ChIP-seq tracks for CTCF and H3K27ac and ATAC-seq tracks corresponding to the *BCL11B* (c) and *TLX3* (d) loci in T cell, Jurkat and Loucy T-ALL cells are shown below. Chromosomal breakpoints mapped by this study are marked by vertical lines with case number indicated. Lower: Expression of *BCL11B* and *TLX3* in each sample. *BCL11B-TLX3* translocations lead to ectopic TLX3 expressions in cases 93 and 117.
