## Supplementary Figure 4 for "3D Genome Analysis Identifies Enhancer Hijacking Mechanism for High-Risk Factors in Human T-Lineage Acute Lymphoblastic Leukemia"

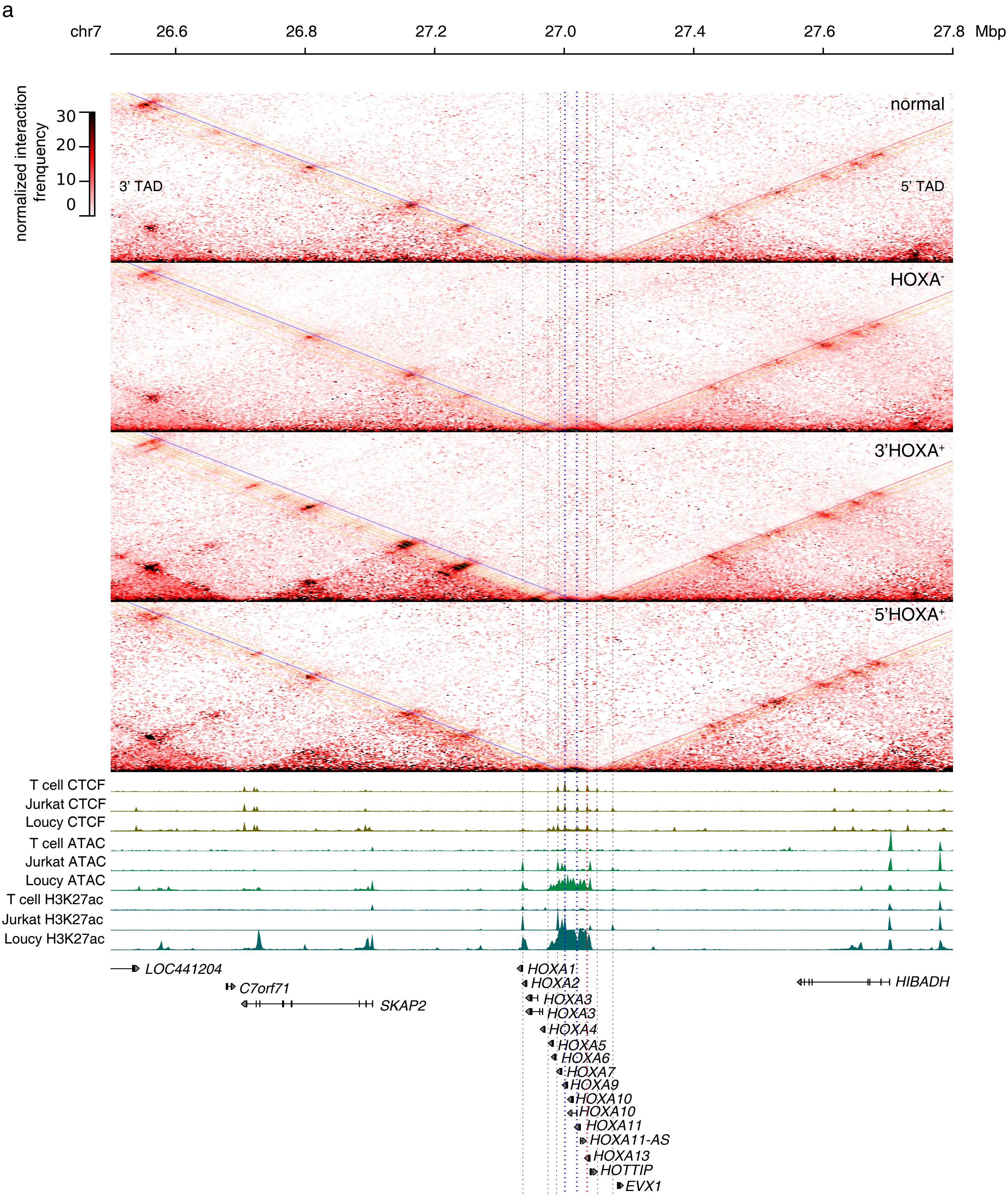

a
chr7
26.6
26.8
27.2
27.0
27.4
27.6
27.8
Mbp
normalized interaction frenquency
30
20
10
0
3' TAD
5' TAD
normal
HOXA-
3'HOXA+
5'HOXA+
T cell CTCF
Jurkat CTCF
Loucy CTCF
T cell ATAC
Jurkat ATAC
Loucy ATAC
T cell H3K27ac
Jurkat H3K27ac
Loucy H3K27ac
LOC441204
C7orf71
SKAP2
HOXA1
HOXA2
HOXA3
HOXA3
HOXA4
HOXA5
HOXA6
HOXA7
HOXA9
HOXA10
HOXA10
HOXA11
HOXA11-AS
HOXA13
HOTTIP
EVX1
HIBADH

b

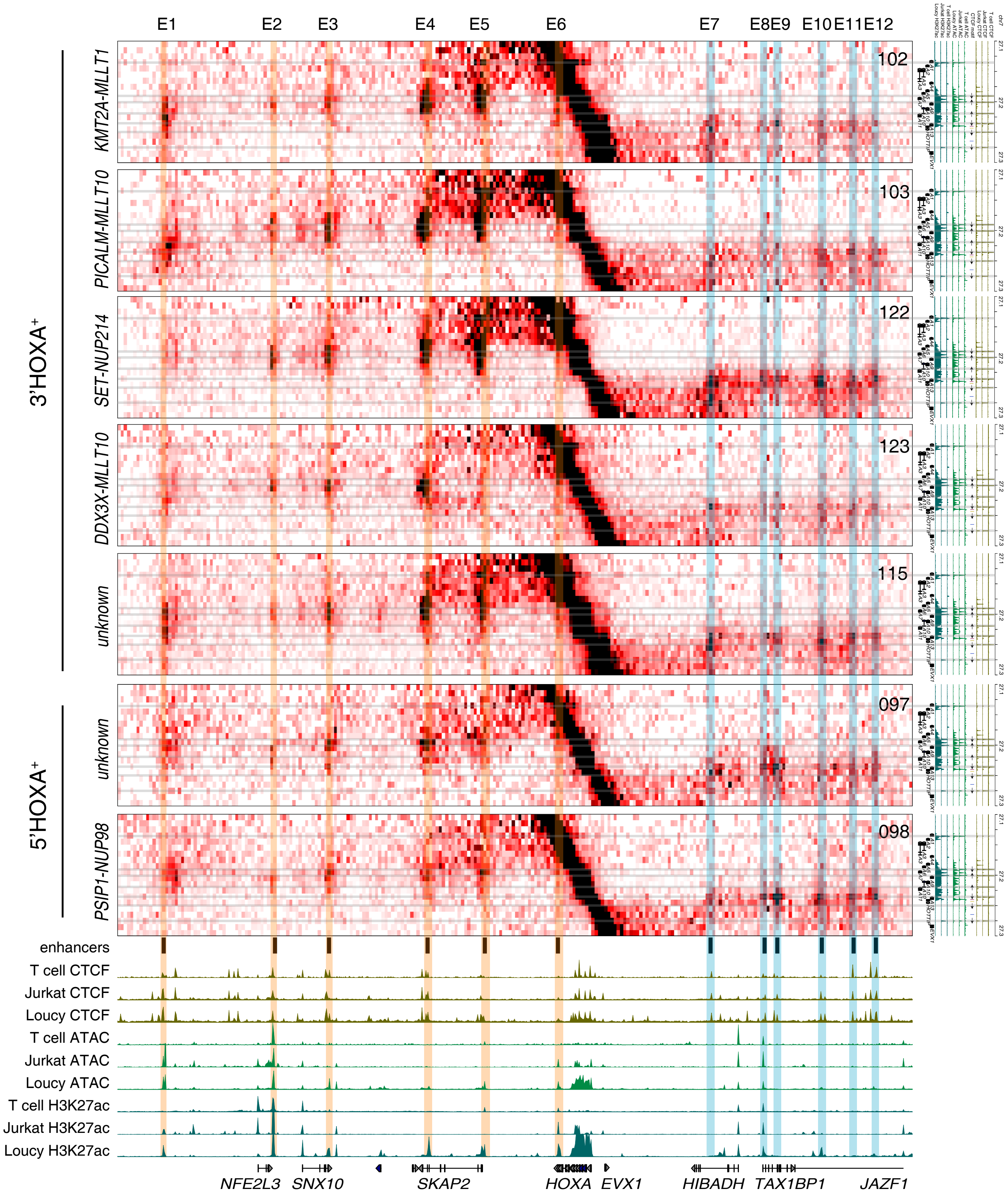
E1
E2
E3
E4
E5
E6
E7
E8
E9
E10
E11
E12
3'HOXA+
5'HOXA+
KMT2A-MLLT1
PICALM-MLLT10
SET-NUP214
DDX3X-MLLT10
unknown
unknown
PSIP1-NUP98
102
103
122
123
115
097
098
enhancers
T cell CTCF
Jurkat CTCF
Loucy CTCF
T cell ATAC
Jurkat ATAC
Loucy ATAC
T cell H3K27ac
Jurkat H3K27ac
Loucy H3K27ac
NFE2L3
SNX10
SKAP2
HOXA
EVX1
HIBADH
TAX1BP1
JAZF1

c

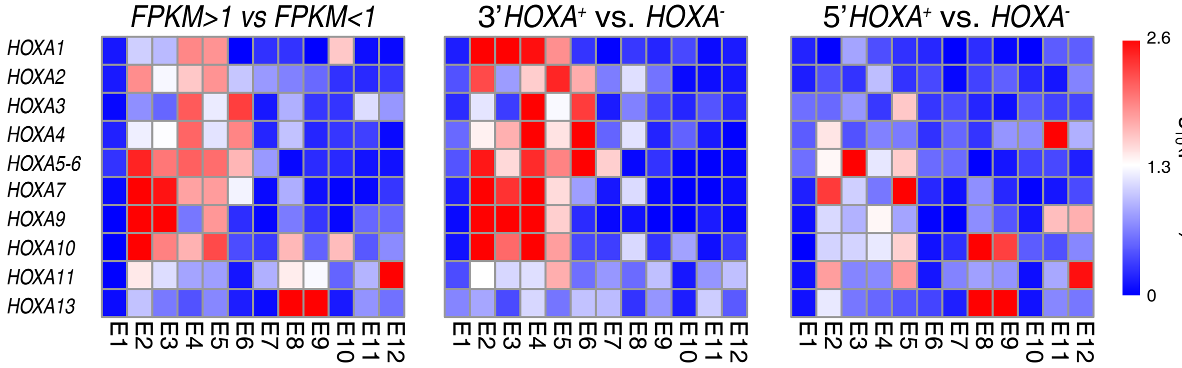
FPKM>1 vs FPKM<1
3'HOXA+ vs. HOXA-
5'HOXA+ vs. HOXA-
HOXA1
HOXA2
HOXA3
HOXA4
HOXA5-6
HOXA7
HOXA9
HOXA10
HOXA11
HOXA13
E1
E2
E3
E4
E5
E6
E7
E8
E9
E10
E11
E12
2.6
1.3
0
-log10(p-value)

**Supplementary Fig. 4 Chromatin interaction profile and expression patterns of the *HOXA* cluster in T-ALLs.** (a) Hi-C heatmaps show the average interaction intensity of normal T cells, $HOXA^-$, 3'$HOXA^+$ and 5'$HOXA^+$ cases in chr7: 25,750,000-28,250,000 (hg19), which includes *HOXA* gene cluster and its 3' and 5' TADs. Vertical black dotted lines mark the CTCF binding sites near the 3' TAD and 5' TAD boundaries; red line marks the 3' boundary of 5' TAD in all samples while blue lines mark the two 5' boundaries of 3' TAD among different samples. ChIP-seq tracks for CTCF and H3K27ac and ATAC-seq tracks corresponding to the *HOXA* cluster in T cell, Jurkat and Loucy T-ALL cells are shown below. (b) Individual Hi-C contact maps between genomic region chr7: 25,750,000-28,250,000 and chr7:27,100,000-27,300,000 for 3'$HOXA^+$ and 5'$HOXA^+$ cases. The main enhancers are highlighted with orange in the 3' TAD and blue in the 5' TAD. The CTCF sites are heighted with grey horizontal bars. (c) Heatmap showing statistical test conducted by comparing interactome of each HOXA gene between its expression as FPKM > 1 and FPKM < 1 (left), 3'HOXA+ and HOXA- cases (middle), 5'HOXA+ and HOXA- cases (right). *P* value is calculated with t-test.
