## Supplementary Figure 5 for "3D Genome Analysis Identifies Enhancer Hijacking Mechanism for High-Risk Factors in Human T-Lineage Acute Lymphoblastic Leukemia"

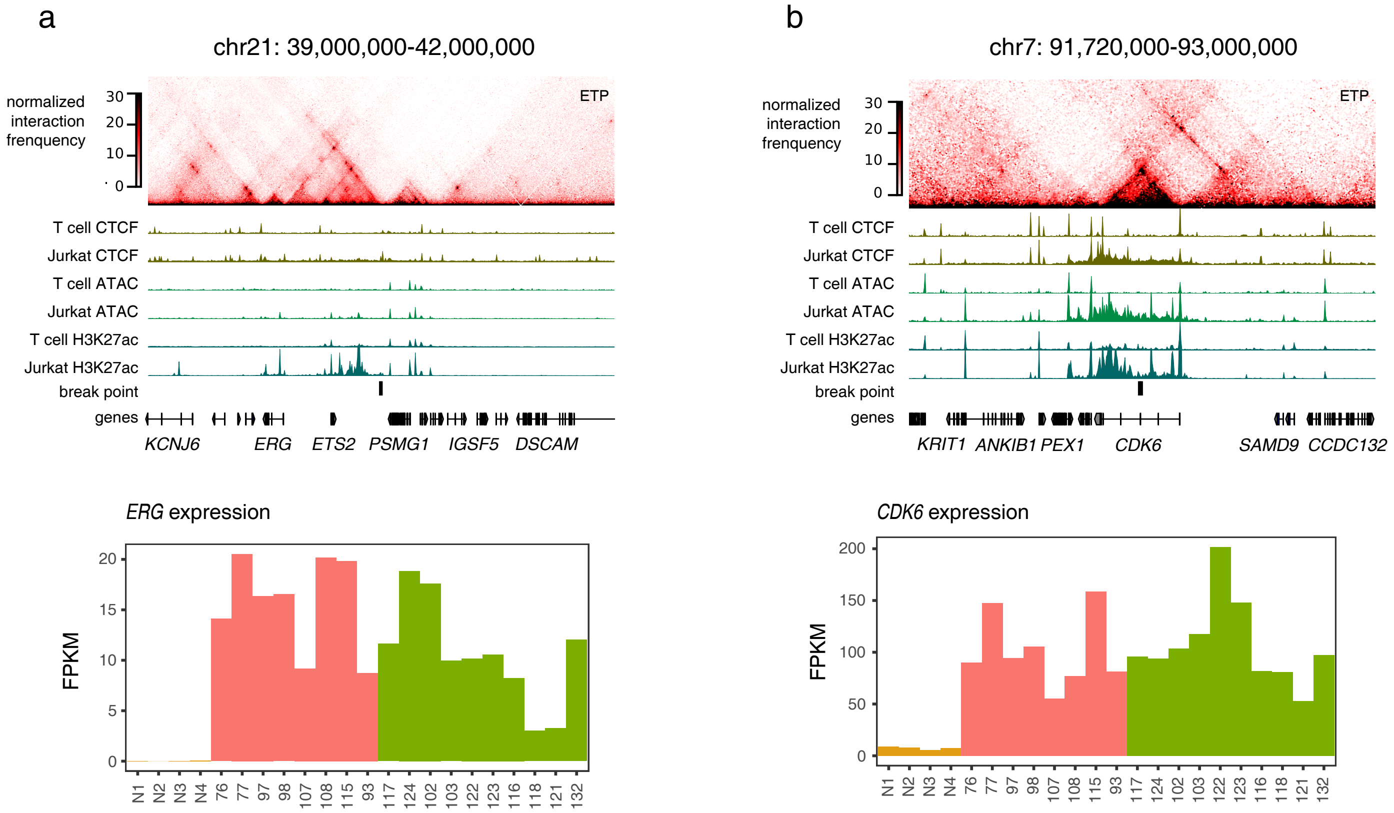


**Supplementary Fig. 5 Translocation-mediated enhancer hijacks and ectopic *HOXA* gene expressions in T-ALLs.** (a) and (b) Upper: Hi-C contact maps for TADs enclosing the genomic loci of *ERG* (a) and *CDK6* (b) genes. Lower: The expression levels of *ERG* and *CDK6* genes in each sample.
