## Supplementary Figure 6 for "3D Genome Analysis Identifies Enhancer Hijacking Mechanism for High-Risk Factors in Human T-Lineage Acute Lymphoblastic Leukemia"

a

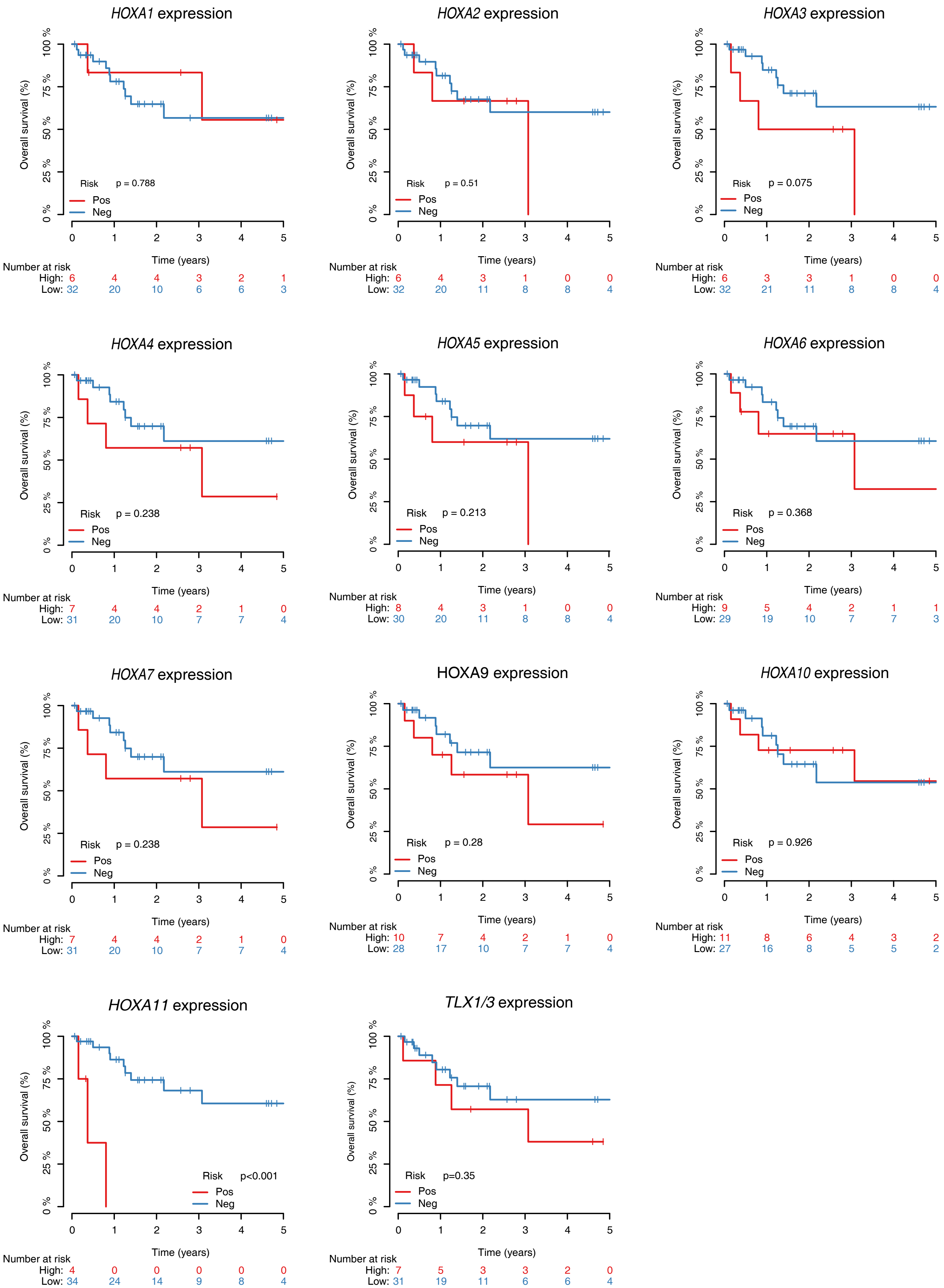

HOXA1 expression
Overall survival (%)
100 %
75 %
50 %
25 %
0 %
Risk p = 0.788
Pos
Neg
Time (years)
Number at risk
High: 6 4 4 3 2 1
Low: 32 20 10 6 6 3
HOXA2 expression
Risk p = 0.51
Pos
Neg
Number at risk
High: 6 4 3 1 0 0
Low: 32 20 11 8 8 4
HOXA3 expression
Risk p = 0.075
Pos
Neg
Number at risk
High: 6 3 3 1 0 0
Low: 32 21 11 8 8 4
HOXA4 expression
Risk p = 0.238
Pos
Neg
Number at risk
High: 7 4 4 2 1 0
Low: 31 20 10 7 7 4
HOXA5 expression
Risk p = 0.213
Pos
Neg
Number at risk
High: 8 4 3 1 0 0
Low: 30 20 11 8 8 4
HOXA6 expression
Risk p = 0.368
Pos
Neg
Number at risk
High: 9 5 4 2 1 1
Low: 29 19 10 7 7 3
HOXA7 expression
Risk p = 0.238
Pos
Neg
Number at risk
High: 7 4 4 2 1 0
Low: 31 20 10 7 7 4
HOXA9 expression
Risk p = 0.28
Pos
Neg
Number at risk
High: 10 7 4 2 1 0
Low: 28 17 10 7 7 4
HOXA10 expression
Risk p = 0.926
Pos
Neg
Number at risk
High: 11 8 6 4 3 2
Low: 27 16 8 5 5 2
HOXA11 expression
Risk p<0.001
Pos
Neg
Number at risk
High: 4 0 0 0 0 0
Low: 34 24 14 9 8 4
TLX1/3 expression
Risk p=0.35
Pos
Neg
Number at risk
High: 7 5 3 3 2 0
Low: 31 19 11 6 6 4

b

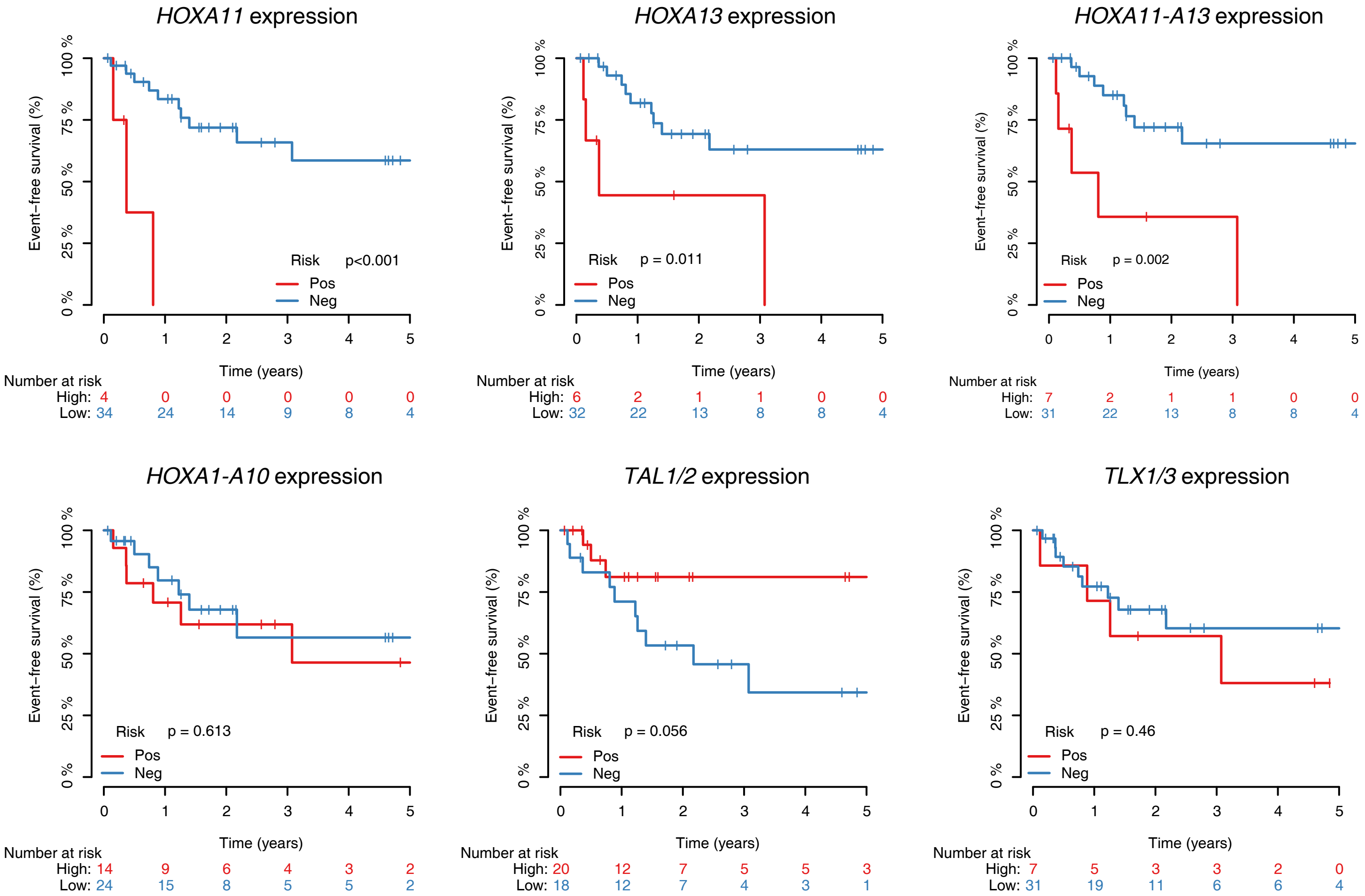

HOXA11 expression
HOXA13 expression
HOXA11-A13 expression
HOXA1-A10 expression
TAL1/2 expression
TLX1/3 expression
Event−free survival (%)
0 % 25 % 50 % 75 % 100 %
Time (years)
0 1 2 3 4 5
Risk p<0.001
Risk p = 0.011
Risk p = 0.002
Risk p = 0.613
Risk p = 0.056
Risk p = 0.46
Pos
Neg
Number at risk
High: 4 0 0 0 0 0
Low: 34 24 14 9 8 4
High: 6 2 1 1 0 0
Low: 32 22 13 8 8 4
High: 7 2 1 1 0 0
Low: 31 22 13 8 8 4
High: 14 9 6 4 3 2
Low: 24 15 8 5 5 2
High: 20 12 7 5 5 3
Low: 18 12 7 4 3 1
High: 7 5 3 3 2 0
Low: 31 19 11 6 6 4


c

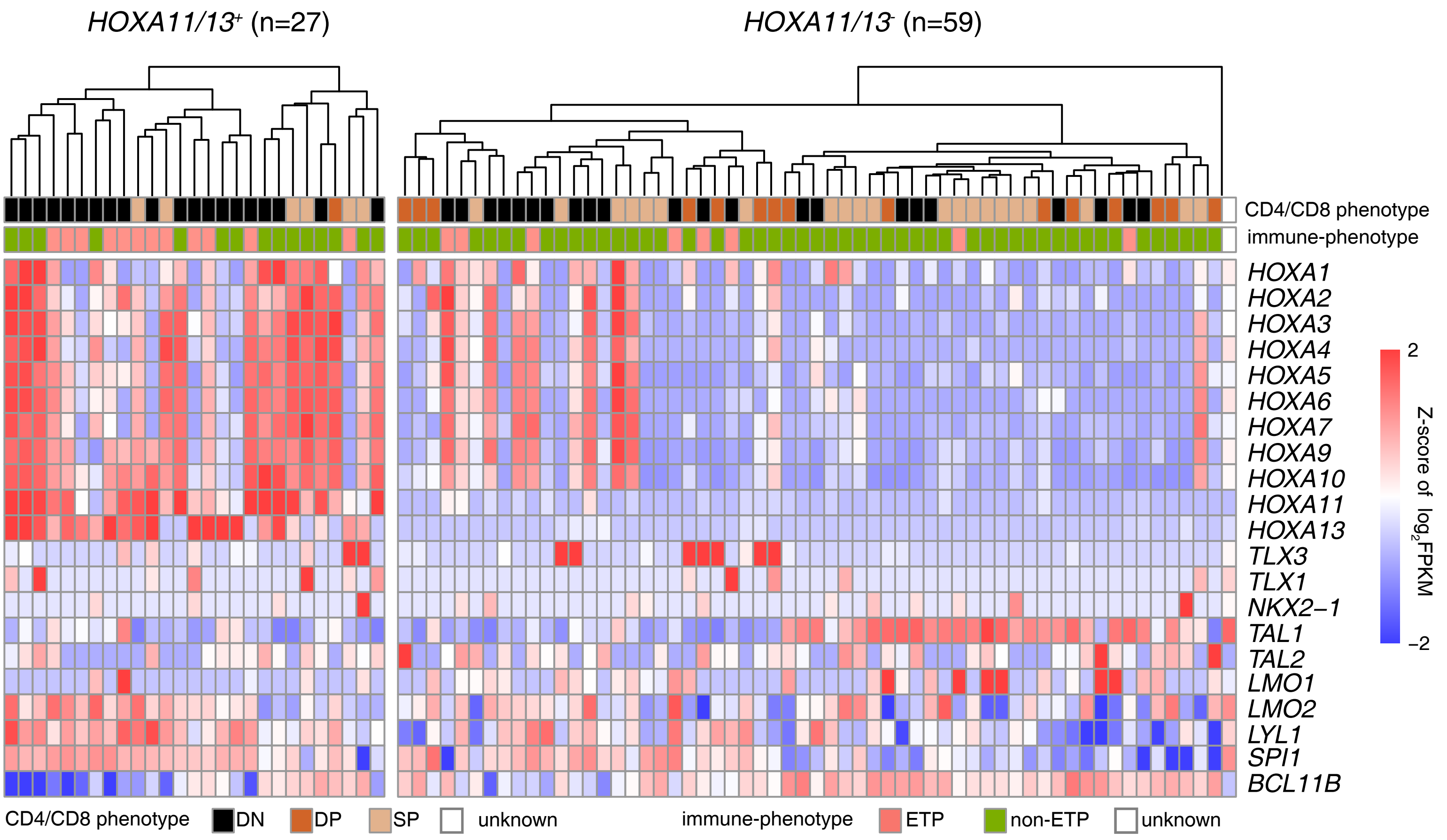

HOXA11/13+ (n=27)
HOXA11/13- (n=59)
CD4/CD8 phenotype
immune-phenotype
HOXA1
HOXA2
HOXA3
HOXA4
HOXA5
HOXA6
HOXA7
HOXA9
HOXA10
HOXA11
HOXA13
TLX3
TLX1
NKX2−1
TAL1
TAL2
LMO1
LMO2
LYL1
SPI1
BCL11B
2
−2
Z-score of log2FPKM
CD4/CD8 phenotype DN DP SP unknown
immune-phenotype ETP non-ETP unknown

**Supplementary Fig. 6 Ectopic *HOXA11-A13* expressions are correlated with poor outcomes in pediatric and young adult T-ALLs.** (a) Kaplan-Meier overall survival curves of pediatric and young adult patients with (red) or without (blue) *HOXA1* to *HOXA11* and *TLX1/TLX3* expressions. (b) Kaplan-Meier event free survival curves of pediatric and young adult patients with (red) or without (blue) *HOXA11* or *HOXA13* expression, *HOXA11-A13* expression, *HOXA* expression excluding HOXA11 or A13 expression (*HOXA1-A10*), T-ALL subtypes specific oncogenic transcription factor *TAL1/TAL2* and *TLX1/TLX3* expression. *P* values are calculated by the log-rank test. (c) A heatmap shows the profile of leukemogenic transcription factors expression in a larger T-ALL cohort (86 cases). Unsupervised clustering of $HOXA11/13^{+}$ and $HOXA11/13^{-}$ subgroups demonstrates preferential expression of early T cell development associated transcription factors *LMO2*, *LYN1* and *SPI1* and association of DN and ETP phenotypes in HOXA11/13+ subgroup.
